# Inhibiting EZH2 complements steroid effects in Duchenne muscular dystrophy

**DOI:** 10.1101/2024.08.22.609220

**Authors:** Eun Young Jeon, Yejin Kwak, Hyeji Kang, Se Young Jin, Soojin Park, Ryeo Gyeong Kim, Dayoung Ko, Jae-Kyung Won, Anna Cho, Inkyung Jung, Chul-Hwan Lee, Jeongbin Park, Hyun-Young Kim, Jong-Hee Chae, Murim Choi

## Abstract

Duchenne muscular dystrophy (DMD) is a devastating X-linked disorder caused by mutations in the dystrophin gene. Despite recent advances in understanding the disease etiology and applying emerging treatment methodologies, glucocorticoid derivatives remain the only general therapeutic option that can slow disease development. However, the precise molecular mechanism of glucocorticoid action remains unclear, and there is still need for additional remedies to complement the treatment. Here, using single-nucleus RNA-sequencing and spatial transcriptome analyses of human and mouse muscles, we investigated pathogenic features in DMD patients and palliative effects of glucocorticoids. Our approach further illuminated the importance of proliferating satellite cells, and revealed increased activity of a signal transduction pathway involving EZH2 in the patient cells. Subsequent administration of EZH2 inhibitors to *Dmd* mutant mice resulted in improved muscle phenotype through maintaining the immune-suppressing effect but overriding the muscle weakness and fibrogenic effects exerted by glucocorticoids. Our analysis reveals pathogenic mechanisms that can be readily targeted by extant therapeutic options for DMD.

**Teaser:** A survey of DMD tissues in human and mouse suggests EZH2 as a critical factor in DMD satellite cells; its inhibition resulted in better prognosis.

## Introduction

Duchenne muscular dystrophy (DMD) is a severe, progressive, and muscle-wasting X-linked genetic disorder that affects about 1 in 5,000 males (*1*).

Mutations in the dystrophin gene are responsible for both DMD and Becker muscular dystrophy (BMD), a relatively less severe and heterogeneous form of dystrophinopathy (*2*). The severe symptoms of DMD and relative ease of access to the affected tissue made it one of the first target diseases for emerging cell- and gene-based therapeutic approaches. However, attempts involving viral transfer of the *DMD* gene or gene activation machinery via the CRISPR system have yielded unsatisfactory outcomes (*3–5*). Therefore, while studies have demonstrated gene editing therapies, there remain challenges in validating the clinical benefits and controlling potential off-target effects that need to be addressed before clinical application (*6*).

At present, the standard of care for DMD still relies on glucocorticoids (*e.g*., deflazacort and prednisone), mitigating the disease progression by reducing inflammation-induced muscle damage, which minimizes muscle strength loss (*7–9*). Indeed, treatment with deflazacort delays the onset of cardiomyopathy in DMD patients until the age of 18 and increases patient survival by over 5–15 years (*10*). However, the precise action of deflazacort in DMD muscle tissue is still unclear, and its long-term administration has well-documented side effects such as obesity, behavioral changes, short stature, and osteopenia (*11*). There have been a few attempts to treat DMD using chemical compound; for example, cyclosporine is administered as an immunosuppressant (*12, 13*), suramin attenuates cardiac dysfunction in a DMD mouse model (*14*), and forskolin has improved muscle performance and regenerative capacities in a DMD rat model (*15*). However, while these experimental treatments show promise, it remains crucial to continue pursuing for more effective and safely approved drugs for DMD; especially, a therapeutic that complements deflazacort function would be of great benefit to DMD patients.

Satellite cells constitute a heterogenous population of stem cells that play indispensable roles in the development, preservation, and regenerative processes of skeletal muscles (*16*). These cells can be distinguished by expression of the transcription factor paired-box 7 (PAX7), which has the critical role of enforcing the myogenic program within them (*17*). In DMD, the absence of dystrophin expression leads to impaired regeneration and exacerbating muscle wasting (*18*). In addition to PAX7, polycomb repressive complex 2 (PRC2) has a crucial role in regulating stem-like processes in satellite cells; in cultured skeletal muscle cells, it regulates the cell cycle by controlling proliferation through methylation of histone 3 lysine 27 (H3K27me3). Especially, elevated expression of its histone methyltransferase subunit Ezh2 in cultured mouse cells has been shown to inhibited muscle differentiation (*19*). Therefore, perturbation of EZH2 in satellite cells may provide a new opportunity for influencing myocyte differentiation in skeletal muscle.

To further understand the molecular mechanism of deflazacort in DMD and to suggest additional therapeutic options for the disease, we performed a comparative analysis of human DMD and BMD patients and *Dmd* mutant mice with or without deflazacort treatment. This study design intended selective utilization of human and mouse systems in modeling DMD and steroid treatment. Subsequently, we reveal disease-specific features in DMD muscle tissues and demonstrate molecular mechanisms underlying the beneficial effects of deflazacort, as well as new potential therapeutic strategies for using EZH2 inhibitors, GSK126 and Tazemetostat, to improve muscle phenotype in combination with glucocorticoids. The beneficial effect was found not only in satellite cells, but also in other cell types, such as immune cells and fibro/adipogenic progenitors (FAPs). Ultimately, our results elucidate a compensatory mechanism of stem cell stimulation on the established effect of deflazacort.

## Results

### Cellular profiles of muscle tissues in human cases and a DMD mouse model

Single-nucleus RNA-seq (snRNA-seq) was performed on fresh frozen muscle cells obtained from three male individuals with DMD, three male individuals with BMD, and five healthy control subjects (Methods; Fig. 1A and fig. S1). All participants were under the age of 17. Of the patients with BMD, two exhibited deletion mutations (p.Glu2147_Gln2171del and exon 30-42 del), while the third had a stop codon mutation (p.Ser1273X) in dystrophin gene. Among the DMD cases, two had stop codon mutations (p.Arg195X and p.Gln423X), and one patient had a frameshift insertion mutation (p.His3299Glnfs*15) (table S1).

Following snRNA-seq of these muscle samples, quality filtering was performed and 60,886 nuclei were retained for subsequent analyses. We captured 11 main cell types, including type I and II muscle cells, FAPs, satellite cells, and myeloid cells (Fig. 1B). Notably, patients with BMD and DMD exhibited expansion of FAP and myeloid lineages, concomitant with a decline in muscle cell populations (Fig. 1C, D, fig. S2). The proportion of immune cells in BMD did not show a significant difference from controls.

**Fig. 1.**
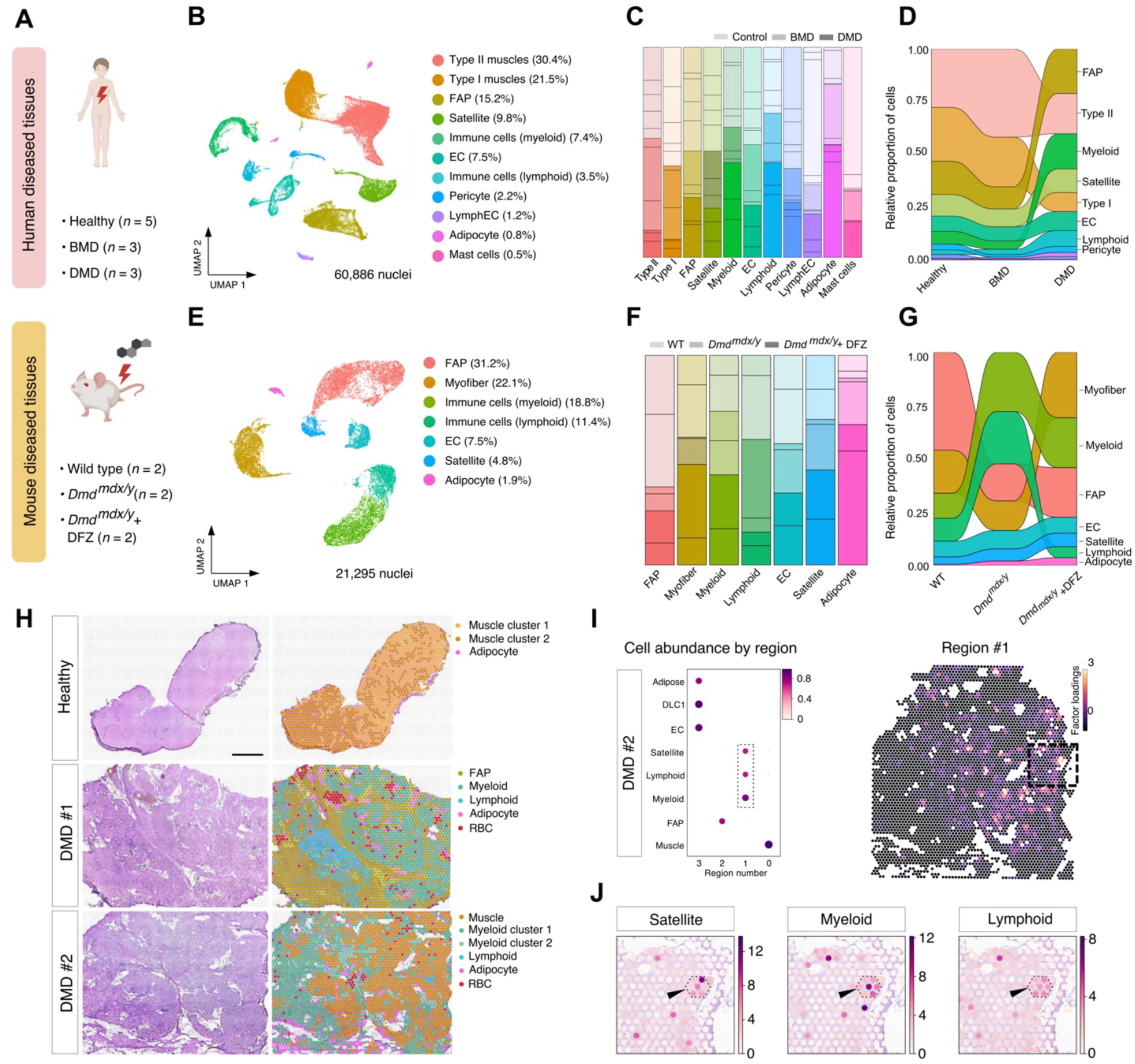
Profiles of human and mouse DMD muscle tissues. (**A**) Human and mouse tissues used in the study. DFZ denotes deflazacort. (**B, E**) UMAP visualization of 60,886 nuclei from human (**B**) and 21,295 nuclei from mouse (**E**) tissues. FAP, fibro-adipogenic progenitors; EC, endothelial cells; LymphEC, lymphatic endothelial cells. (**C, F**) Cell type proportions in human (**C**) and mouse (**F**) samples. (**D, G)** Changes in cell type proportion by disease state in human (**D**) and mouse (**G**) samples. (**H**) Cell type mapping on Visium slides of human DMD and healthy muscle sections. Scale bar = 1 mm. (**I**) Dot plot illustrating co-occurrence of spatially-resolved cell types in regions defined by cell2location (*69*). The dotted rectangle emphasizes the co-occurrence of satellite cells and immune cells in region #1 (left). Spatial heatmap showing the location of region #1 (right). (**J**) Magnified view of the dotted rectangle in (I) (right), emphasizing colocalized expression of satellite, myeloid, and lymphoid markers in the DMD #2 sample.

We also performed snRNA-seq on fresh frozen muscle cells from two male wild-type DBA/2J mice, two male D2.B10-*DMD^mdx^*/J (or *mdx*) mice, and two male *mdx* mice administered with deflazacort (Methods, Fig. 1A and fig. S1). The two mice in each genotype group is consisted of a 7- and a 28-week-old mouse, as *mdx* mice show a notable decline in skeletal muscle function as early as seven weeks and in cardiac function by 28 weeks (*20*). Seven distinct cell types were identified from 21,295 nuclei (Fig. 1E). Like human patient muscles, *mdx* muscles exhibited a decrease in muscle cells and an increase in myeloid and lymphoid cell subpopulations (Fig. 1F, G). In *mdx* animals subjected to deflazacort treatment, noticeable reduction of myeloid cell frequencies and increase of muscle cells were evident. Inflammatory myeloid cell population marked by *Cd44* and *Itgam* genes drove this change (fig. S2). In the comparative analysis between 7- and 28-week-old mice, we repeatedly observed a notable increase of Wnt signaling gene expression. The hallmark was upregulation of *Tcf7l2*, a transcription factor for Wnt signaling, target genes of which were confirmed to have increased expression in seven-week-old *mdx* mice as assayed by SCENIC (fig. S3). This observation was not seen in wild type mice or in *mdx* mice given deflazacort.

Concurrently with the snRNA-seq profiling, spatial transcriptomics was conducted on a subset of two DMD patients and one healthy samples using the Visium method (Fig. 1H and fig. S4). The control subject showed mainly muscle-specific expression profiles, aligning with the observed cellular composition. However, the two slides derived from DMD patients exhibited substantial augmentation of myeloid cell expression profiles, indicating a pronounced shift in the cellular landscape. Cell type co-occurrence analysis further demonstrated an increase in the co-occurrence of satellite cells, myeloid cells, and lymphoid cells, suggesting local interactions among these cells in the DMD muscle (Fig. 1I, J).

### Effects of deflazacort in humans and a mouse DMD model

To investigate the molecular consequences of deflazacort treatment in humans and a mouse models of DMD, we conducted a comprehensive investigation of differentially expressed genes (DEGs). In myeloid cells, a cluster of guanine nucleotide exchange factors (GEFs) encompassing *DOCK2*, *DOCK8*, and *DOCK10* displayed significant upregulation across BMD, DMD, and *mdx* mice when compared to corresponding controls (Fig. 2A, B), implying upregulation of RhoA-mediated actin remodeling, cell migration, and subsequent immune activation (*21, 22*). Administration of deflazacort resulted in a decreased expression of these genes. Binding sites for NR3C1, a dedicated nuclear receptor for glucocorticoid, were identified near *DOCK2*, *DOCK8*, and *DOCK10* loci, suggesting that these genes can be direct downstream factors of deflazacort (Fig. 2C, fig. S5).

**Fig. 2.**
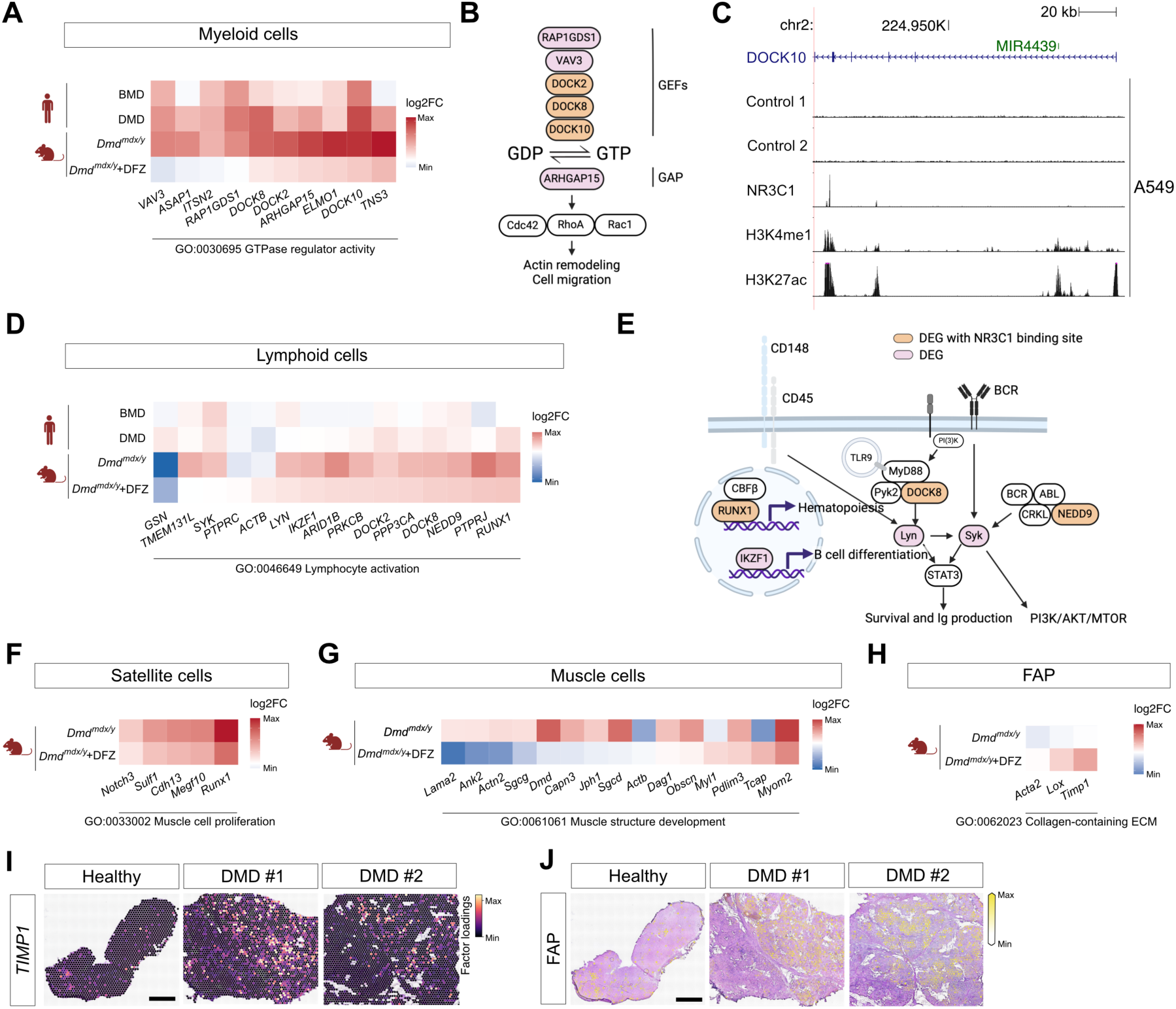
Altered gene expression in human and mouse tissues with *DMD* mutation and deflazacort treatment. (**A**, **D**, **F**, **G, H**) Heatmaps of DEGs in each cell type by species and disease status. **A**: myeloid cells, **D**: lymphoid cells, **F**: satellite cells, **G**: muscle cells, and **H**: FAP cells. (**B**, **E**) Schematic representation of the roles of DEGs in corresponding heatmaps. **B**: actin remodeling and cell migration genes in myeloid cells and **E**: B cell differentiation, survival and Ig production genes in lymphoid cells. (**C**) Diagram of the exemplary NR3C1 ChIP-seq peak locus on *DOCK10,* a DEG found in the myeloid cells. (**I**) *TIMP1* expression in FAP clusters. (**J**) Visualization of FAP clusters by cell2location mapping on Visium slides of human patient muscle sections. Scale bars = 1 mm.

Similarly, we investigated DEGs within lymphoid cells (Fig. 2D). Genes linked to hematopoiesis, B cell differentiation, survival, Ig production, and the PI3K/AKT/MTOR pathway exhibited increased expression across BMD, DMD and *mdx* lymphocytes (Fig. 2E). Upon deflazacort administration, expression of these genes was attenuated (Fig. 2D).

In contrast to the discernible improvements seen in myeloid and lymphoid cells following deflazacort intervention, no enhancement was observed within satellite, muscle, and FAP clusters (Fig. 2F, G, H). Indeed, genes related to regulation of satellite cell growth and muscle regeneration were depleted (Fig. 2F) (*23–26*), and muscle structural genes (Fig. 2G) displayed decreased expression following deflazacort administration. The concomitant elevated presence of pro-fibrotic FAP markers (Fig. 2H, I) suggests a potential avenue for intervention as FAP cells are capable of differentiating towards either adipogenic or myofibroblast lineages (Fig. 2J) (*27, 28*). Therefore, although deflazacort positively affects immune cells through controlling inflammatory signaling pathways, it also exacerbates muscle structure deficits and fibrosis in DMD muscles.

### *EZH2* is expressed in proliferating satellites

Since deflazacort mainly exerts its beneficial effects in muscle through immune modulation, we sought to identify a distinct pathway that can provide additional, complementary benefit in DMD treatment. First, we examined variations in gene expression across the trajectory of muscle differentiation from satellite cells to myocytes among control, BMD and DMD muscles. Figure 3A focuses on the muscle structure development and muscle cell differentiation pathways that were specifically upregulated in controls compared to BMD and DMD muscles.

**Fig. 3.**
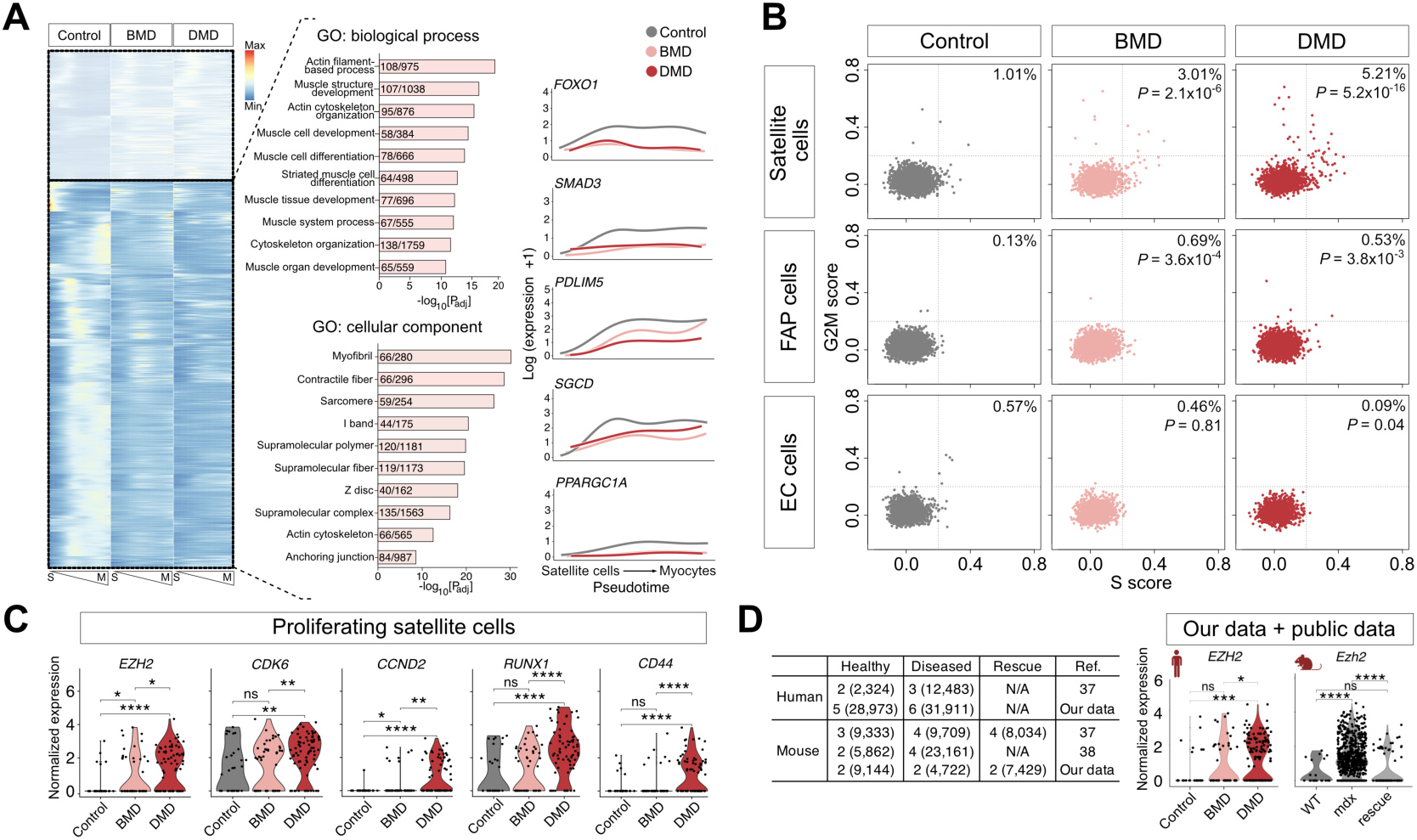
Proliferating satellite cells in DMD patients lead to increased signal transduction in a pathway involving *EZH2* and cell cycle progressors. (**A**) Heatmap of genes whose expressions vary over pseudo-temporal ordering from satellite cells (S) to myocytes (M) (left). GO terms enriched among these genes (middle). Numbers in bar graphs indicate numbers of input genes/genes in annotation. Plots of log-transformed counts and the fitted values of control, BMD, and DMD patient cells (right). (**B**) Scatterplots of cell cycle scores in satellite, FAP, and EC cells by disease status. Cells were considered positive for cell proliferation if they harbor positive scores for either G2M or S. (**C**) Violin plots depicting DEGs in proliferating satellite cells. (**D**) Table displaying the number of samples and corresponding cell counts (in parentheses) included in our and previous studies (left). Violin plots of both human *EZH2* and mouse *Ezh2* expression in our and public data. ns denotes *P*-value > 0.05, * ≤ 0.05, ** ≤ 0.01, *** ≤ 0.001, **** ≤ 0.0001.

The results of this analysis led us to focus on satellite cells, as they are the source of myocytes (*29, 30*) and proliferate in response to a various kinds of muscle damages (*31–33*). First, we assigned each cell type a quantitative score for cell proliferation based on expression of G2/M and S phase markers (Methods; Fig. 3B). Notably, satellite cells included a group of actively proliferative cells within samples originating from both BMD patients (3.01%; *P* = 2.1 x 10^-6^) and DMD patients (5.21%; *P* = 5.2 x 10^-16^), demonstrating elevation of proliferative activity in compensation for damaged muscle tissues. This observation was specific to satellite cells (Fig. 3B and fig. S6). Similar scoring of cell types was performed for markers of apoptotic and necrotic cell death, but found no substantial differences between healthy and patient cells (fig. S6).

Next, we examined genes showing differential expression profiles in proliferating satellite cells across control, BMD, and DMD muscles. This revealed a group of genes enriched for cell cycle (GO:1903047; *P_adj_* = 9.6 x 10^-15^) (Fig. 3C). Among these, *EZH2* stood out for being known to play a key role in regulating gene expression in muscle cells, and for drugs targeting its gene products being actively used for cancers (*34, 35*). The differential expression of *EZH2* in both DMD patients and *mdx* mice was corroborated through data integration with publicly available datasets (*36, 37*) (Fig. 3D). Therefore, proliferating satellite cells mark increased activity of a signal transduction pathway that engages EZH2 and proteins associated with cell cycle progression in the cells of muscular dystrophy patients.

### Administration of an EZH2 inhibitor improves muscle phenotype in mice

Downregulation of *Ezh2* is known to cause muscle gene activation and myoblast differentiation (*38*), while its increased expression inhibits muscle differentiation *in vitro* (*19*). Therefore, we sought to test if administration of an EZH2 inhibitor elicits an improvement in muscle phenotype in DMD through increased muscle cell differentiation. EZH2 inhibitors, GSK126 and Tazemetostat, were administrated with or without deflazacort to *mdx* mice (Fig. 4A). Representative hematoxylin and eosin (H&E) staining images representing each treatment group’s quadriceps muscles are presented in Fig. 4B; this staining revealed features characteristic of dystrophinopathy. Especially, *Dmd* mice treated with EZH2 inhibitors exhibited improvement in dystrophinopathy in comparison to vehicle-treated mutant mice, in a degree similar or better compared to deflazacort-treated mice (Fig. 4C-E, fig. S7). When relative areas of collagen were compared, *Dmd* mice treated with EZH2 inhibitors exhibited improvement in dystrophinopathy in comparison to vehicle-treated mutant mice, in a degree similar to deflazacort-treated mice (Fig. 4E). Grip strength was significantly increased in mice given Tazemetostat with or without deflazacort compared to deflazacort-only mice (Fig. 4F).

**Fig. 4.**
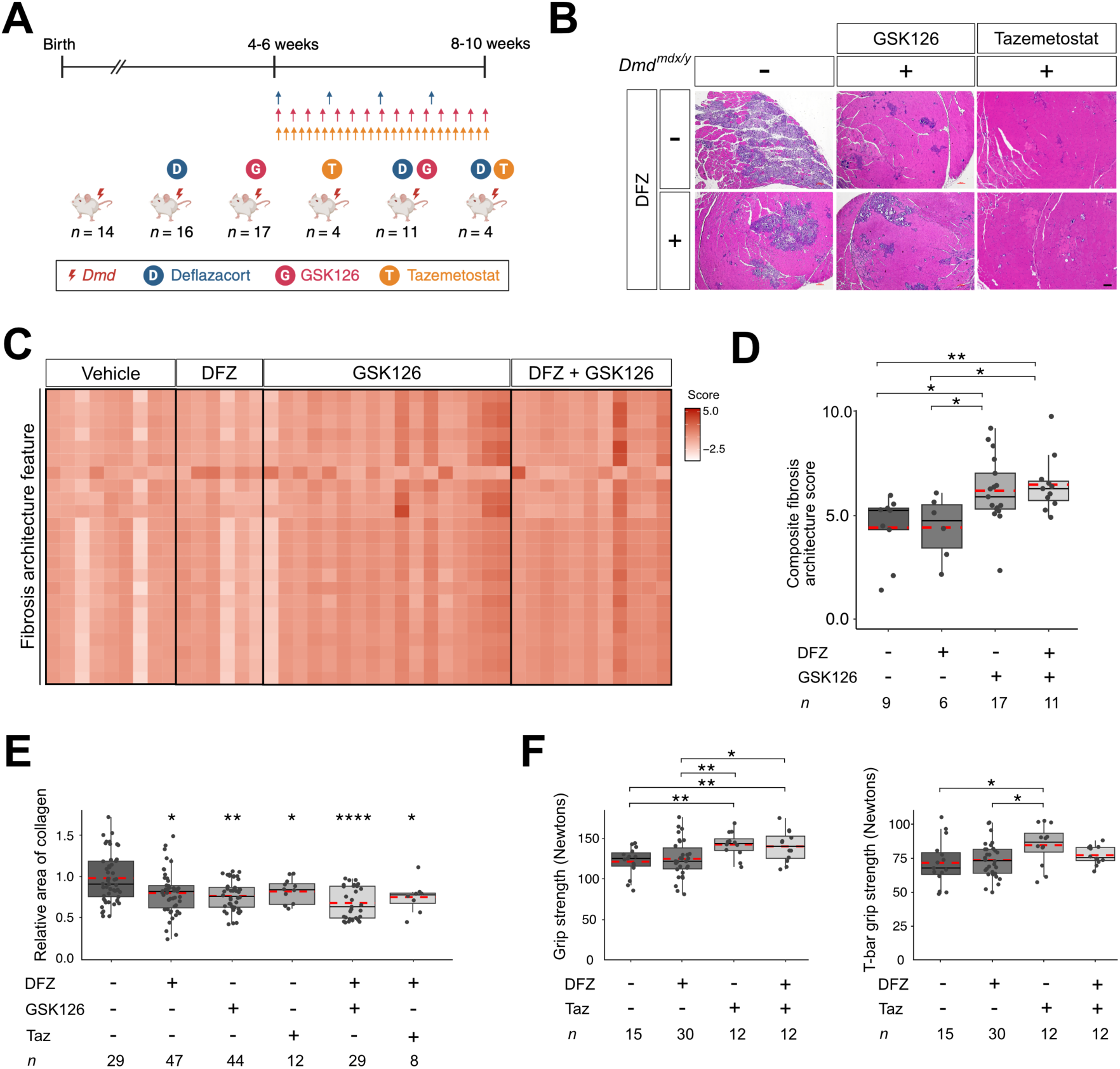
Treatment with EZH2 inhibitors improved muscle phenotype in *mdx* mice. (**A**) Schematic representation of the experimental setup involving mouse drug administration. (**B**) H&E staining of muscle tissues from the 8- to 10-week-old mice after injection of deflazacort and/or GSK126 and/or Tazemetostat (Taz). Scale bar = 100 µm. (**C**) Heatmap showing quantification of Masson Trichrome images for fibrosis architecture features from mice with or without GSK126 treatment. Feature names (row) are listed separately in Data S2. (**D**) Composite fibrosis scores calculated on the basis of (**C**). The number indicates the number of mice used. (**E**) Boxplot showing quantification of area of collagen relative to untreated *mdx* in mice with or without drug treatment based on Masson’s trichrome stained images. The number indicates the number of slides used. (**F**) Boxplots showing grip strength (left) and T-bar grip strength (right) of 8- to 10-week-old mice after injection of deflazacort and/or Tazemetostat. Red dotted lines indicate the mean. The number represents the triplication of the initial count of mice used.

### EZH2 inhibition compensates negative effects of deflazacort in *mdx* mice

To understand the potential consequences of EZH2 inhibition, we performed snRNA-seq on muscle samples from mice given various drug treatments. Following quality control measures, a total of 37,464 nuclei were retained for subsequent analyses (Fig. 5A). Distinct patterns emerged within the cellular landscape, wherein myeloid and lymphoid lineage cells displayed the most pronounced expansion in untreated *mdx* mice, while muscle cells exhibited expansion in *mdx* mice given GSK126 (Fig. 5B). Cell-cycle scoring of myeloid lineage cells revealed a notable variance, with untreated *mdx* mice manifesting the highest proliferative status (*P* < 2.2 x 10^-16^; Fig. 5C). Differential expression analyses of cytokine- and inflammation-related genes uncovered upregulation in untreated *mdx* mice compared to counterparts administered deflazacort and/or GSK126 (Fig. 5D). Especially, the pro-fibrotic genes in FAP cells illustrated in Figure 2H were downregulated in mice that received GSK126 (Fig. 5E).

**Fig. 5.**
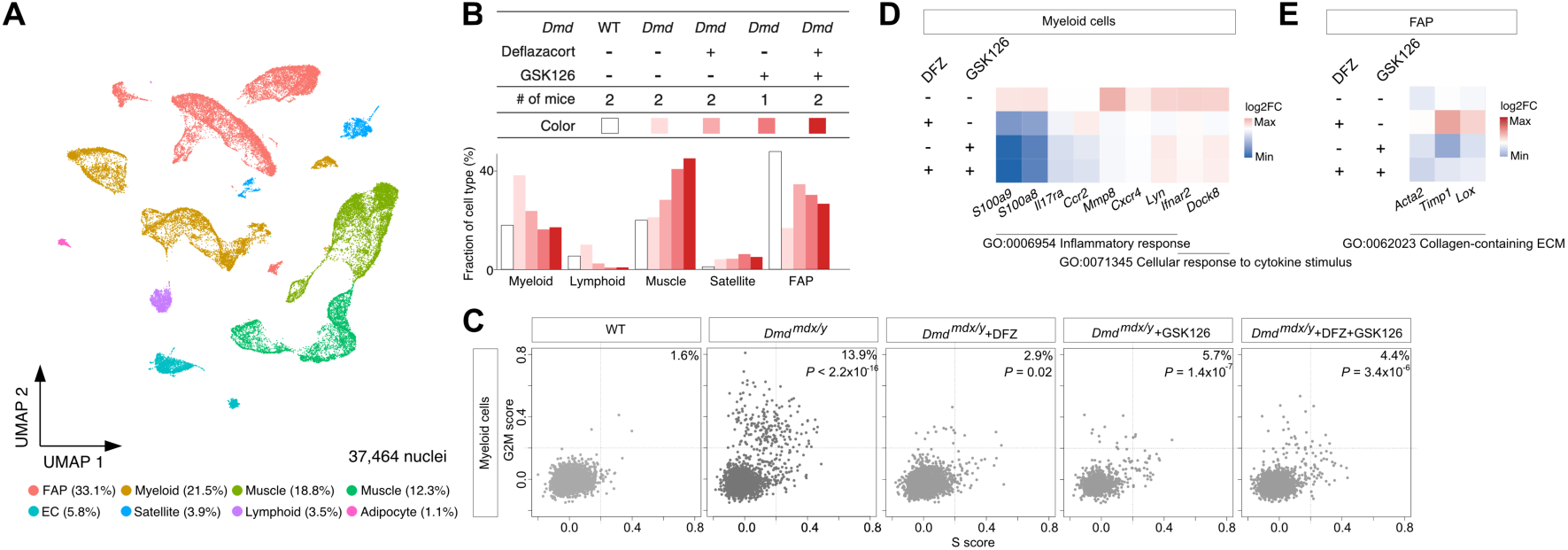
EZH2 inhibitor maintains the immune suppressing effect and overrides the increased fibrosis induced by deflazacort. (**A**) UMAP visualization of 37,464 nuclei from muscle tissues of wild-type or *mdx* mice with drug treatments. (**B**) Cell type proportion by treatment status. (**C**) Scatterplots of cell cycle scores in myeloid cells by treatment status. (**D**) Heatmap of the DEGs in myeloid cells by treatment status. (**E**) Heatmap of the DEGs in FAP cells shown in Fig. 2H by treatment status.

Next, we focused on the beneficial effect of EZH2 inhibition on the myocyte differentiation process. Leveraging single-cell regulatory network inference and clustering analyses, we observed *mdx* mice treated with GSK126 showing activation of *Myog*, a transcription factor pivotal to muscle differentiation, within satellite cells (Fig. 6A, B). In contrast, untreated counterparts did not show *Myog* activation (fig. S8). Figure 6C presents the computed t-distributed stochastic neighbor embedding (t-SNE) visualization of all cells based on the *Pax7* (a pan-satellite cell marker) and *Myog* activity in *mdx* mice given GSK126. Genes associated with muscle structural elements, illustrated in Figure 2G, also demonstrated elevated expression in *mdx* mice treated with GSK126, signifying a potential beneficial effect on muscle architecture (Fig. 6D). To explain this unexpected effect, we queried EZH2-binding sites in muscle cells using ENCODE project data. This revealed a group of genes that were both differentially expressed in proliferating satellite cells and bound by EZH2 to be enriched for cell polarity-related genes (Fig. 6E, F), suggesting a possible restoration of muscle stem cell polarity due to EZH2 inhibition. Binding sites for EZH2 were identified near the *JAM3* locus, one of the genes that were differentially expressed in patients’ proliferating satellite cells and also were bound by EZH2 (Fig. 6F). Indeed, a heatmap of the differentiation trajectory analysis revealed disrupted signatures in *mdx* mice, specifically an overall increase in cell cycle and proliferation markers (Fig. 6G). However, following administration of deflazacort and GSK126, the gene expression pattern resembled that of the wild type. The same pattern persists when GSK126 is given alone (fig. S9).

**Fig. 6.**
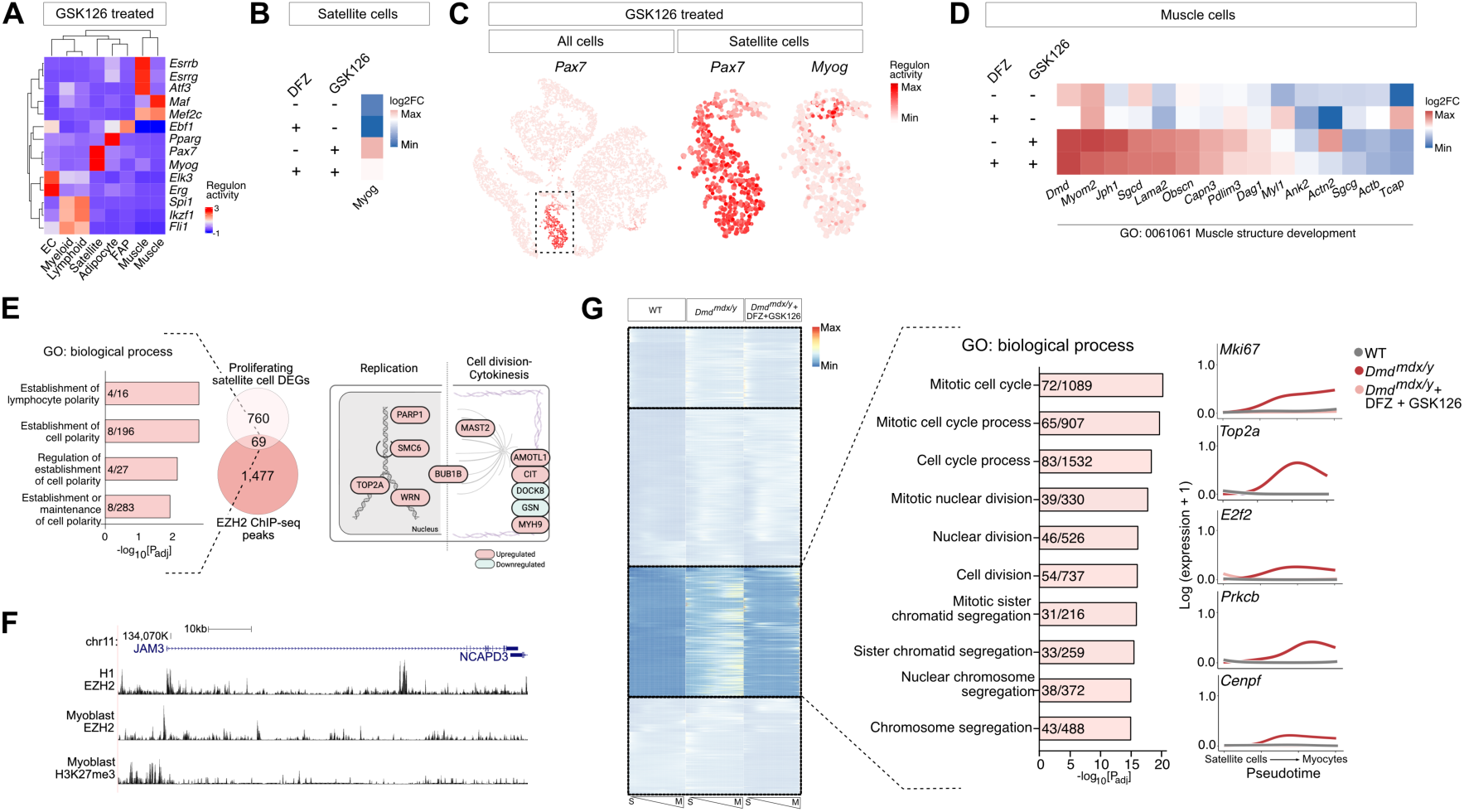
EZH2 inhibitor overrides the muscle weakness effect exerted by deflazacort through stimulating muscle differentiation. (**A**) Heatmap of differentially enriched regulators for each cell type in GSK126-injected mice. (**B**) Heatmap of *Myog* expression in satellite cells by treatment status. (**C**) t-SNE visualization of *Pax7* and *Myog* transcription factor activities in cells of GSK126-injected mice. (**D**) Heatmap of the muscle cell DEGs shown in Fig. 2G by treatment status. (**E**) Number of genes differentially expressed in the proliferating satellite cells and bound by EZH2 (left). Numbers in bar graphs indicate numbers of input genes/genes in annotation. Schematic representation of the roles of overlapping genes that are engaged in cell polarity group (right). (**F**) Diagram of the exemplary EZH2 ChIP-seq peak locus on *JAM3,* a DEG found in the proliferating satellite cells and bound by EZH2. (**G**) Heatmap of genes whose expression varies over pseudo-temporal ordering from satellite cells to the differentiated myocyte state (left). Numbers in bar graphs indicate input genes/genes in annotation. Plots of log-transformed counts and the fitted values of wild type, *mdx* mice, and *mdx* mice with deflazacort and GSK126 injection (right).

## Discussion

DMD is both the most prevalent and the most severe form of muscular dystrophy, typically resulting in a life expectancy limited to the early 20s (*39, 40*). Steroids still remain as the standard of care despite known side effects such as obesity and osteopenia (*41*); development of additional treatments thus remains imperative.

Using snRNA-seq to capture the transcriptomes of individual muscle cells from human patients and model mice, we have uncovered a notable expansion of *EZH2* expression within proliferating satellite cells. Furthermore, we have demonstrated that inhibition of EZH2 has the potential to mitigate immune signals and ameliorate the phenotypic manifestations of DMD.

Currently, four antisense oligonucleotides (ASOs) that are designed to induce skipping of exon 45, 51, or 53 of dystrophin have been granted conditional approval by the Food and Drug Administration (FDA) (*1, 41, 42*). However, the ASO method is specific to particular *DMD* mutations and therefore shows limited applicability. Recently, the FDA also approved a minigene therapy exclusively for the treatment of ambulatory DMD pediatric patients within the specific age range of 4-5 years (*3*). However, a clinical benefit such as enhanced motor function has not been confirmed, and the FDA has mandated the company to conduct a comprehensive clinical study to validate the drug’s clinical advantages as a condition for maintaining approval (*4*). Moreover, even patients successfully treated with these approaches will still be administered steroids. Therefore, there is still critical need for a therapeutic method that can be applied on a broad range of DMD patients and complement steroids.

A few recent studies have examined the pathophysiology of DMD muscles at the single-cell level. Scripture-Adams *et al.* analyzed skeletal muscle cellular diversity in human patients and *mdx* mice with and without exon skipping therapy (*36*).

Another study observed increased expression of FAP cells, consistent with our findings, and upregulation of plasminogen activator inhibitor-1 (*PAI1*) in dystrophic endothelial cells (*37*). In the skeletal muscle of mice with *DMD* exon 51 deletion, another study identified distinctive myonuclear subtypes within dystrophin myofibers and explored transcriptional pathways associated with degeneration and regeneration in DMD (*43*). Consistent with the previous reports, we observed both DMD patients and *mdx* mice displaying reduced myocytes and increased immune cells, which imbalance was partly reverted in mice by the administration of deflazacort. The profiles of BMD muscle were closer to normal than those of DMD (Fig. 1D), reflecting the milder symptoms of BMD patients. It would be of interest to further investigate the genetic and phenotypic correlation of *DMD* mutations and disease severity (*44*). Remarkably, our study uncovered a co-localization of satellite cells and immune cells in the inflamed muscle tissue of DMD patients (Fig. 1I-J, fig. S10). This close proximity of satellite cells and immune cells may possibly account for the increased expression of *EZH2* and increased inflammation observed in DMD patients. Although steroids are commonly used in the clinic, the particular molecular cascades they alter have remained elusive. In this study, we found that deflazacort treatment reduced actin remodeling and inflammatory signals in immune cells through NR3C1 binding elements, suggesting a possible mode of action of steroids in DMD treatment (Fig. 2).

Previous studies have observed cellular consequences of reduced EZH2 function in various biological contexts. In cancers, *EZH2* knockdown induces cell cycle arrest and impairs *in vitro* migration and invasion (*45–47*), while depletion of *Ezh2* results in diminished activation of macrophages and microglia (*48*). Here we further demonstrated molecular mechanisms underlying the immunosuppressive effects of Ezh2 inhibition. Inflammatory monocyte-derived macrophages were known to play a critical role in DMD pathogenesis, and CCR2 deficiency helped restore the macrophage polarization balance by preventing an excessive shift towards a proinflammatory phenotype (*49*). *CCR2* was decreased with EZH2 inhibitor treatment (Fig. 5D), suggesting that CCR2 could potentially be targeted therapeutically with EZH2 inhibitor.

Through genome-wide mapping of histone modifications in muscle satellite cells and studying mice lacking *Ezh2* in satellite cells, it was discovered that Ezh2 activity is required for satellite cell proliferation (*50*). In the context of DMD, *EZH2* appears to be a double-edged sword. On one hand, increased presence of EZH2 in satellite cells could be beneficial, potentially serving as a compensatory mechanism to counteract the detrimental effects of DMD through supporting satellite cell proliferation. However, inhibiting EZH2 may encourage satellite cells to differentiate and contribute to muscle repair. Striking the right balance between promoting proliferation and differentiation of satellite cells may optimize therapeutic strategies for DMD patients. A previous study demonstrated that satellite cell-specific ablation of *Ezh2* in mice resulted in reduced muscle mass and impaired regeneration (*51*). This experimental outcome reflects different experimental conditions, with ours being based on pharmacological inhibition in mature muscle. Nevertheless, the study also observed increased expression of myogenic and satellite cell-specific expression, consistent with our findings (*51*). Here, we elucidated the precise role of EZH2 and its implications in DMD treatment. The augmented expression of EZH2 in proliferating satellite cells supports a state of active proliferation and hence hindered muscle differentiation (*19*) (Fig. 3). Administration of EZH2 inhibitors in *mdx* mice reduced fibrosis (Fig. 4). These findings underscore the potential contribution of EZH2 inhibitors to the attenuation of adipogenic or fibrotic regions (Fig. 5E). Furthermore, we observed GSK126 administration bolstering the expression of genes associated with muscle organization through forming a series of differentiating lineages from *Pax7^+^Myog^−^*, *Pax7^+^Myog^+^* and finally the *Pax7^-^Myog^+^* fetal myocytes-like lineage (Fig. 6A-C, fig. S11) (*52*). The beneficial effect was not only found in satellite cells, but also in other cell types.

Recently, Auger *et al.* demonstrated a non-transcriptional pathway that steroid can incur anti-inflammatory effects by reshaping the mitochondrial metabolism of macrophages, leading to increased tricarboxylic acid cycle (TCA) activity and subsequent itaconate synthesis (*53*). Increased TCA activity was also observed in our data with mice given deflazacort alone (fig. S12). However, of these signatures no longer persisted in mice given EZH2 inhibitor. It appears that the inhibition of EZH2 overrides the upregulation of TCA cycle.

Although our findings suggest EZH2 inhibition as a novel therapeutic option that can be used in conjunction with deflazacort, more detailed molecular mechanisms are still needed. The ChIP-seq analysis of proliferating satellite cells might offer insights into the direct targets of EZH2 inhibition. However, such an experiment cannot presently be performed owing to the paucity of human proliferating satellite cells. We anticipate that the advent of new methodologies that require less sample input will help to elucidate these direct targets. In addition to that methodological limitation, it is notable that myofibers in DMD are known to undergo necrotic and apoptotic processes (*54, 55*); however, none of the these signatures were detected in our analysis (fig. S6). Presumably the cells harboring these signatures were filtered out during the quality control step. We aimed to utilize two different species assuming their conserved response to *DMD* mutation. However, there appears to be a noticeable discrepancy in the effect on lymphoid cells (Fig. 2D). Specifically, despite the parallel responses observed in myeloid cells (Fig. 2A), human lymphoid cells exhibited a lesser degree of susceptibility compared to their mouse counterparts.

Collectively, our comparative analysis of human patients and a mouse DMD model have unveiled genes and pathways that exhibit selective downregulation in immune cells and concurrent upregulation in muscle organizational cells. This discovery suggests a novel therapeutic target with promising implications for the treatment of DMD.

## Materials and Methods

### Human participants

Muscle biopsies from patients were performed in accordance with informed written consent from Seoul National Hospital Children’s Hospital institutional review board (IRB)-approved protocols (#1009-030-331). The patients did not take any medication at the time of the biopsy. Muscle samples were taken from the quadriceps femoris and frozen with isopentane cooled in liquid nitrogen as described previously (*56*). Healthy muscle samples were acquired from the quadriceps or abdomen of subjects undergoing surgery for non-muscular symptoms.

### Mouse strains

DBA/2J and DBA/2J-mdx mice (D2.B10-*Dmd^mdx^*/J), hereafter denoted as *mdx* mice, were purchased from the Jackson Laboratory (JAX ID: 013141; Bar Harbor, ME). All experiments were approved by the Institutional Animal Care and Use Committee in Seoul National University Hospital (#20-0216-S1A0) and animals were maintained in a facility accredited by AAALAC International (#001169) in accordance with the Guide for the Care and Use of Laboratory Animals 8^th^ edition, NRC. Muscle samples were taken from the hindlimb and quadriceps and frozen with liquid nitrogen.

### Generating and processing snRNA-seq data

All muscle samples were processed and snRNA-seq data generated by GENINUS (Seoul, Republic of Korea). After the frozen tissue was homogenized and nuclei were counted, the nuclei were isolated using flow cytometry. Single-cell capture, barcoding and library preparation were performed following the 10x Genomics Single cell Chromium 3’ protocols (V3: CG000183). cDNA library quality was determined using an Agilent Bioanalyzer. Paired-end 200 bp reads were generated on an Illumina NovaSeq5000/6000.

### Analysis of snRNA-seq data

Generated FASTQ files were mapped to either human (GRCh38/hg38 pre-mRNA genome) or mouse (mm10) transcriptome references provided by 10x Genomics using Cell Ranger v6.0.0. The output was processed using Seurat v4.0.1 (*57*). We then applied standard cell filtering criteria (nFeature_RNA > 200, nFeature_RNA < 5,000, percent.mt < 5). Nuclei that passed the filtering were normalized and integrated with LIGER v1.0.0 (*58*).

Genes that were differentially expressed between patient and control tissues were defined in each cluster using the FindMarker function of Seurat. SCENIC v1.3.1 was used to infer transcription factor-target relationships (*59*). CellChat was used to infer intercellular communications (*60*). Slingshot was used for cell lineage and pseudotime inference (*61*). TradeSeq was used for trajectory-based differential expression analysis (*62*). Gene set enrichment analysis was performed using the *escape* R package, v1.99.0 (*63*). For mouse immune sub-clustering analysis, data from the Single-Cell Muscle Project was used (*64–67*).

### Gene set enrichment analysis

Cell-cycle scores were assigned using the CellCycleSoring() function in Seurat. Scoring was based on the strategy described by Tirosh *et al.* (*68*). Gene set enrichment analysis was performed using the escape R package v1.99.0 (*63*). Gene sets were derived from the Molecular Signature Database (https://www.gsea-msigdb.org/gsea/msigdb/).

### Generation, processing, and analysis of spatial transcriptomics data

Samples processing and data generation using the Visium platform was done by GENINUS. Experiments were performed using the Visium Spatial Platform 3’ v1 (PN-1000193, PN-1000184, PN-1000215). Sequencing reads from Visium ST (10x Genomics) experiments were first preprocessed with Space Ranger v1.3.1 and mapped to the human reference genome (GRCh38). The count matrices were subsequently analyzed using cell2location v0.1.3 (*69*). To discriminate transcriptionally distinct cell populations with MERFISH, we designed a panel of 300 genes selected based on cluster markers identified from snRNA-seq data.

### ChIP-seq data analysis

ChIP-seq data for NR3C1 (A549 cells: ENCFF638NRS), H3K4me1 (A549 cells: ENCFF040HPO), H3K27ac (A549 cells: ENCFF541LPH), EZH2 (myoblast: ENCFF353VYD; H1 cells: ENCFF109KCQ), and H3K27me3 (myoblast: ENCFF261INX) in the bigwig format were downloaded from the ENCODE project (https://www.encodeproject.org/) and visualized in UCSC Genome Browser (*70*).

ChIPseeker (*71, 72*) was used to annotate ChIP peaks.

### Drug treatment

Deflazacort (SML0123-10MG; Sigma, St. Louis, MO) was formulated as a 0.2 mg/ml suspension in a solution comprised of DMSO (10%), PEG300 (40%), Tween-80 (5%), and saline (45%). This formulation was administered at a dose of 1 mg/kg by intraperitoneal injection once a week for 28 days to four week-old mice. GSK126 (S7061; Selleck chemicals, Houston, TX) was administered to four-week-old mice by intraperitoneal injection once every two days for 28 days at a dose of 50 mg/kg in 20% Captisol (*35, 73*). Tazemetostat (S7128; Selleck Chemicals, Houston, TX) (400 mg/kg, 0.5% CMCNa and 0.1% Tween80 in water) was orally administrated daily for 28 days (*74, 75*).

### Histology and image quantification

After embedding fixed tissues in paraffin, 4-µm sections were stained with Masson’s trichrome stain using the Biognost Masson Trichrome kit (MST-K-500) according to the manufacturer’s protocol and imaged at 40x using a ZEISS Axioscan 7 (ZEISS, Germany). To quantify areas of collagen, Python’s PIL Image and ImageDraw modules were employed to automate the quantification of tissue and collagen regions in Masson’s trichrome histological images. Samples were triplicated. Images were segmented using thresholds and quantified through pixel counting.

For FibroNest (PharmaNest, Princeton, NJ) analysis, the images were cleaned and processed for anomalies such as scanning stripes, image compression artifacts, rinsing artefacts, dusts, and saturated pixels. The digital images were then processed and segmented to allocate the collagen biological marker to a specific channel, as FibroNext selects a region of interest based on collagen fibers. These analyses were blinded to clinical and histological data.

### Behavioral tests

The behavioral assessment was conducted one day prior to tissue collection. Mice were moved to the behavioral assessment room 30 minutes before the test to allow them to acclimate to the new environment. For T-bar grip strength test, the forepaws were stably positioned on the T-bar, and the mouse was pulled gently backward to measure the strength of the forelimb. For grip strength test, the grid was attached securely to the sensor. Both its forepaws and hindpaws were stably positioned. The mouse gently pulled backwards, and while sliding down, the force with which it grips the wire was measured. The tests were repeated three times.

### Statistical analysis

For statistical analysis, R version 4 was used. Statistical analyses for single-cell genomics experiments are described in the preceding subsections. Statistical comparisons between groups were assessed with Fisher’s exact test (Fig. 3B) or *t* tests. A *P*-value of ≤ 0.05 was considered statistically significant.

## Acknowledgments

We express our gratitude to all of the muscle donors and their families for their participation. We thank Young-Yun Kong and Jong-Seol Kang for assistance with muscle dissection and Mathieu Petitjean, Li Chen, and Adi Lightstone at FibroNest for assistance with quantitative AI digital pathology image analysis.

## Author contributions

Conceptualization: MC, J-HC. Methodology: EYJ, YK, HK, SYJ, SP, RGK, J-KW, AC, IJ, C-HL, JP. Investigation: EYJ, YK, HK, AC, C-HL, JP. Visualization: EYJ, MC. Funding acquisition: MC, J-HC, C-HL, IJ. Project administration: EYJ, SP. Supervision: MC, J-HC, JP, C-HL, IJ, AC. Sample acquisition: DK, AC, H-YK, J-HC. Writing – original draft: EYJ, MC. Writing – review & editing: EYJ, YK, HK, SYJ, SP, RGK, DK, J-KW, AC, IJ, C-HL, JP, H-YK, J-HC, MC.

## Data and materials availability

All data are available in the main text or the supplementary materials. All 10x snRNA-seq, 10x Genomics Visium, and Vizgen MERFISH spatial transcriptomics data used in the study are available at the Korea BioData Station with accession numbers <u>temp-grp-2-1711599940117</u>, <u>temp-grp-2-1711600165042</u>, and <u>temp-grp-2-1711600348912</u>, respectively, with minimal restriction posed by the depository.

## Supplementary Materials for

### This PDF file includes

Figs. S1 to S12

Tables S1 to S2

### Other Supplementary Materials for this manuscript include the following

Data S1 to S2

**Fig. S1.**
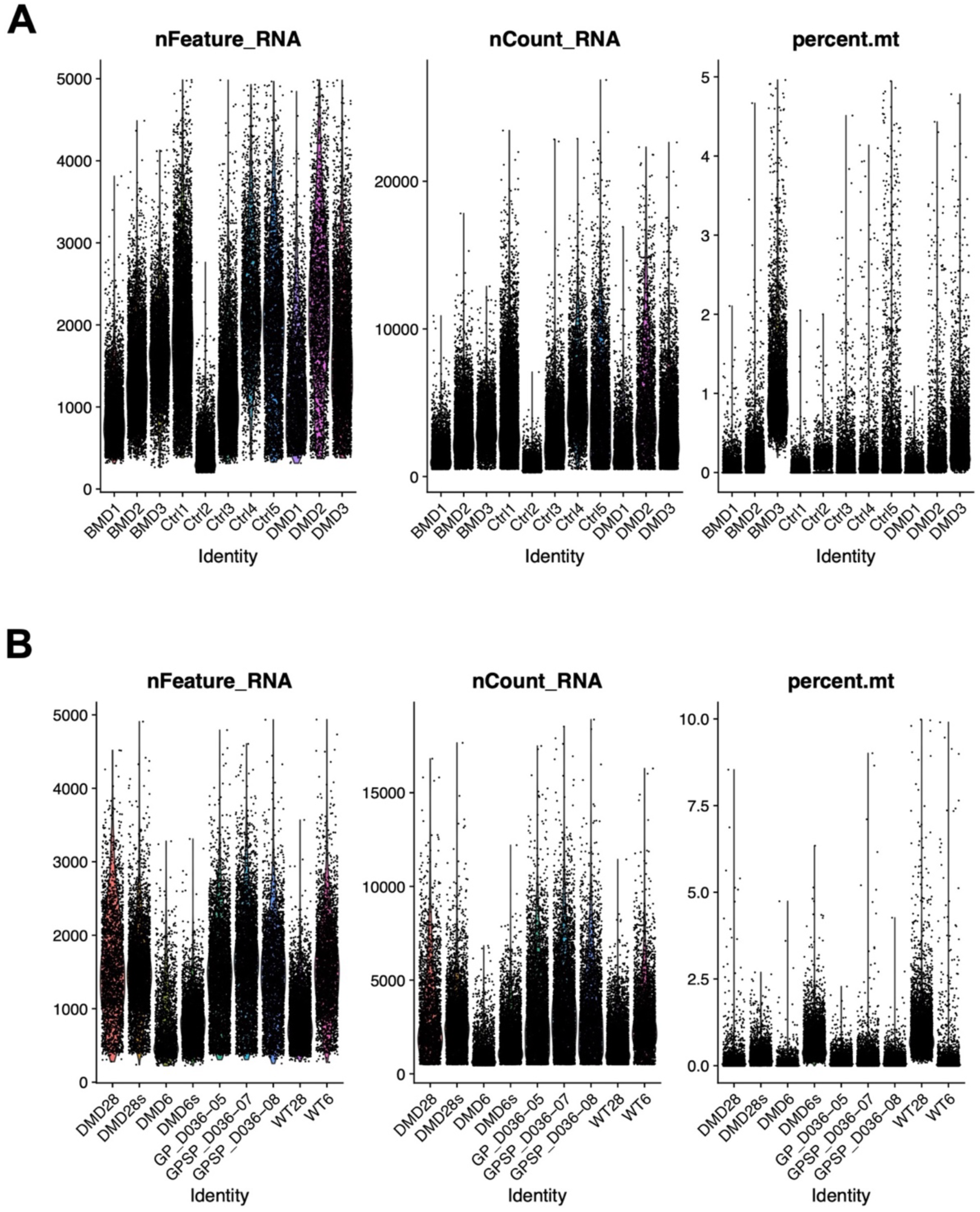
Single nucleus RNA-seq run quality for (A) human and (B) mouse samples.

**Fig. S2.**
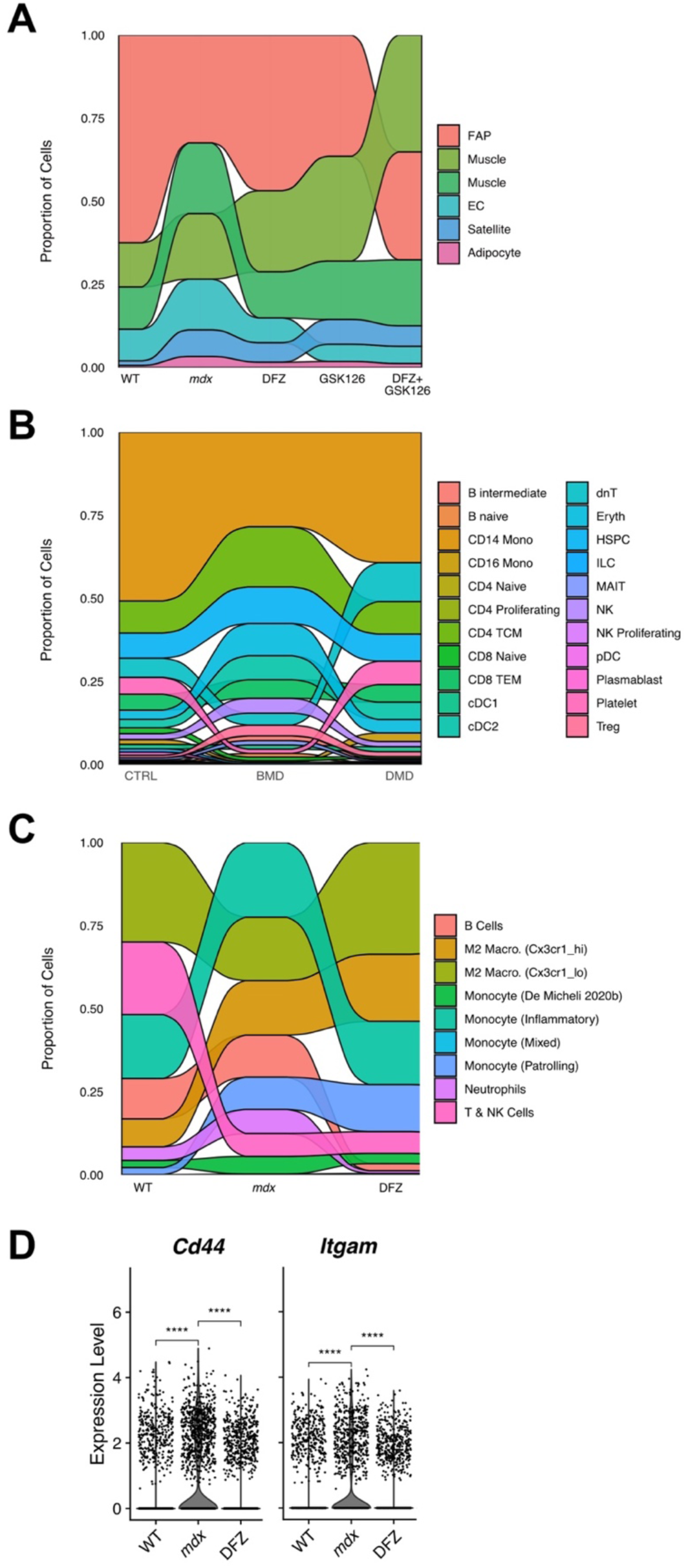
Sub-clustering and differentially expressed genes. **(A)** Mouse data without immune related cells. (**B**) Human data with immune related cells only. (**C**) Mouse data with immune related cells only. (**D**) Differentially expressed genes in the Monocyte (Inflammatory) cluster in (**C**). *** ≤ 0.001.

**Fig. S3.**
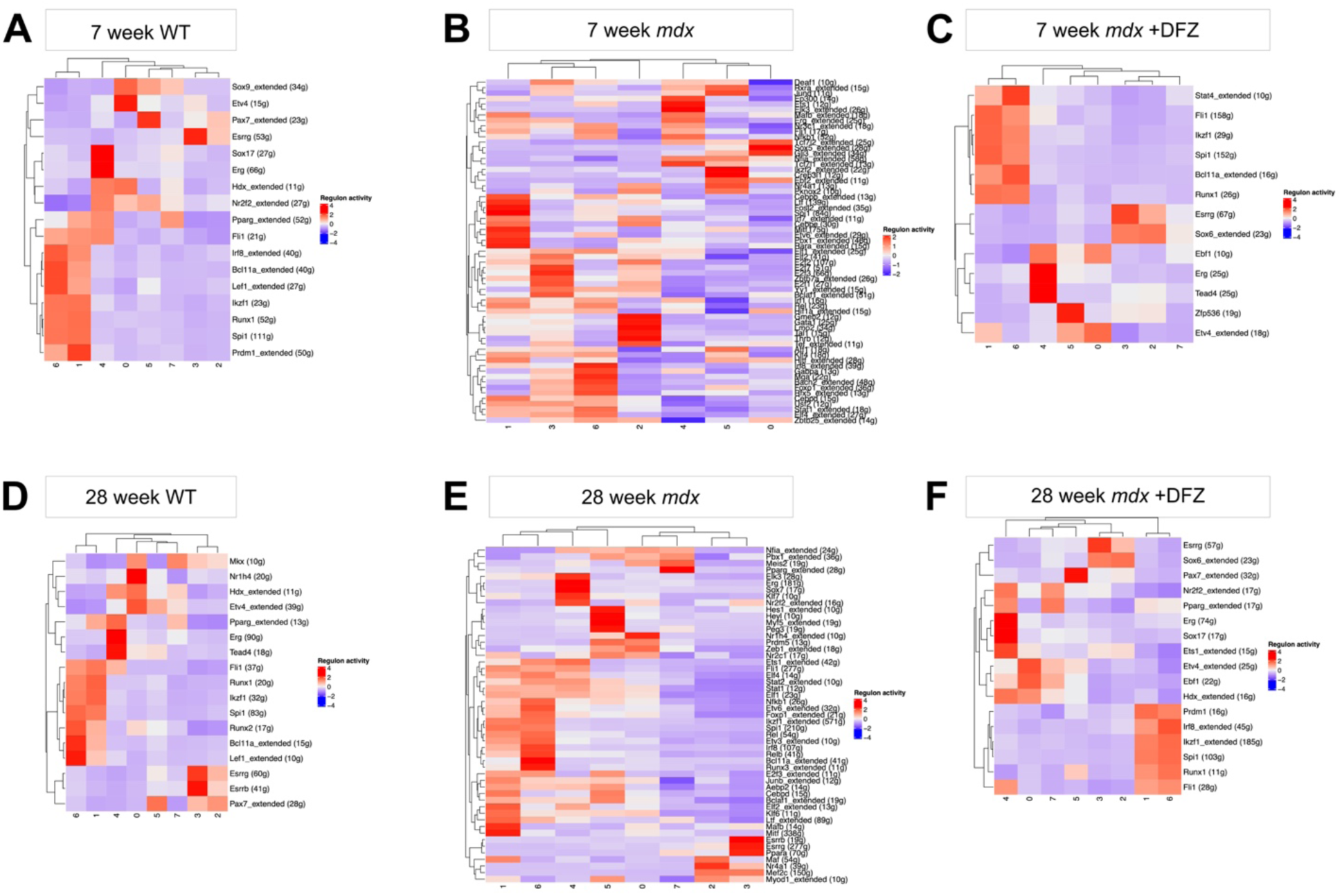
Heatmap of differentially enriched regulators for each cell type. **(A)** Wild type at 7 weeks. (**B**) *mdx* at 7 weeks. (**C**) *mdx* with deflazacort at 7 weeks. (**D**) Wild type at 28 weeks. (**E**) *mdx* at 28 weeks. (**F**) *mdx* with deflazacort at 28 weeks. Numeric labels denote different cell types: 0 - Type II muscle, 1 - Type I muscle, 2 - Fibro-adipogenic progenitors (FAP), 3 - Satellite cells, 4 - Myeloid cells, 5 - Endothelial cells (EC), 6 - Lymphoid cells, 7 - Pericytes, 8 - Lymphatic endothelial cells (lymphEC), 9 - Adipocytes, 10 - Mast cells.

**Fig. S4.**
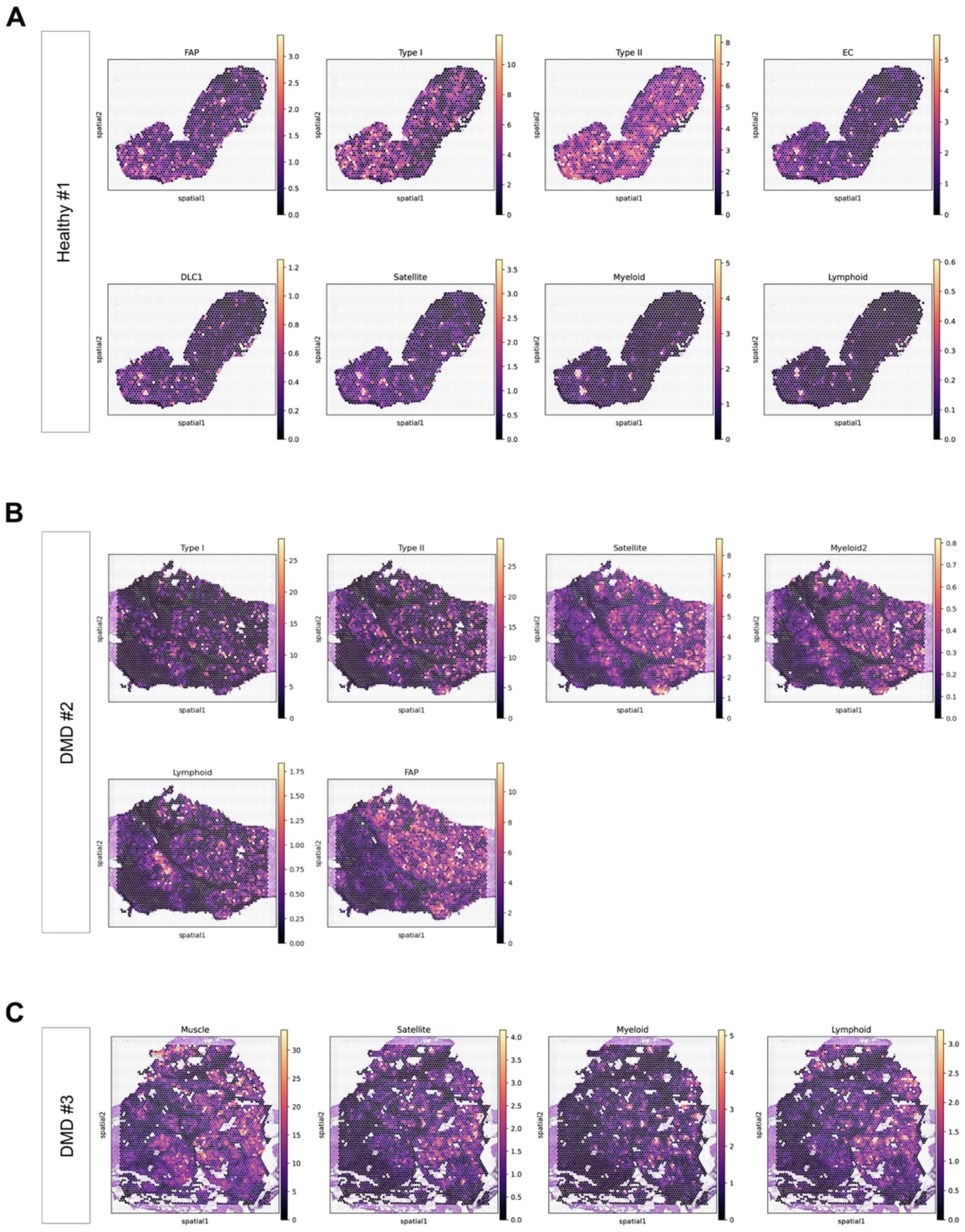
Visium profiles of by cell type by samples. **(A)** A healthy sample. (**B-C**) DMD patient samples.

**Fig. S5.**
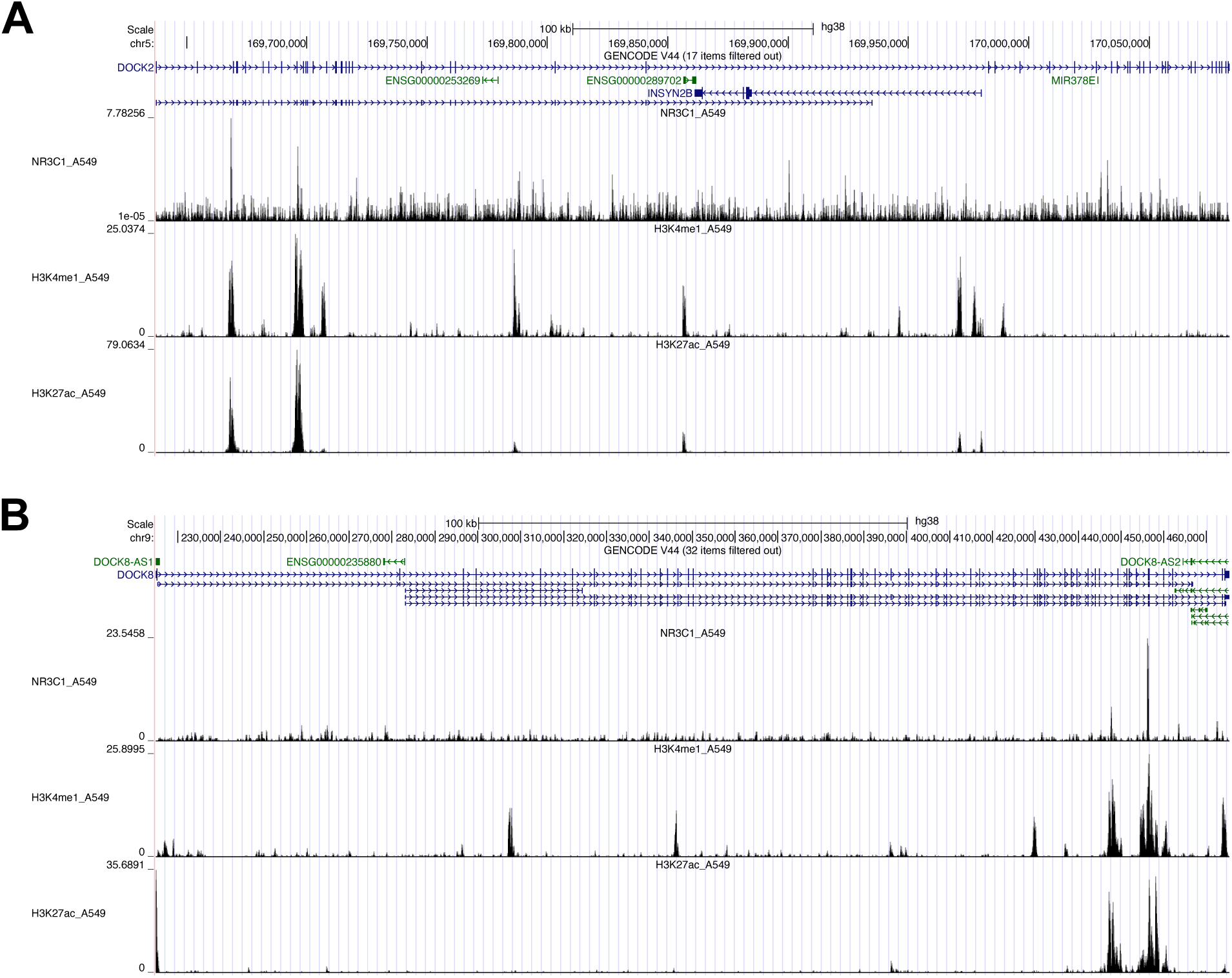
ChIP-seq signals on deflazacort target genes. **(A)** NR3C1 ChIP-seq peak locus on *DOCK2*. (**B**) NR3C1 ChIP-seq peak locus on *DOCK8*.

**Fig. S6.**
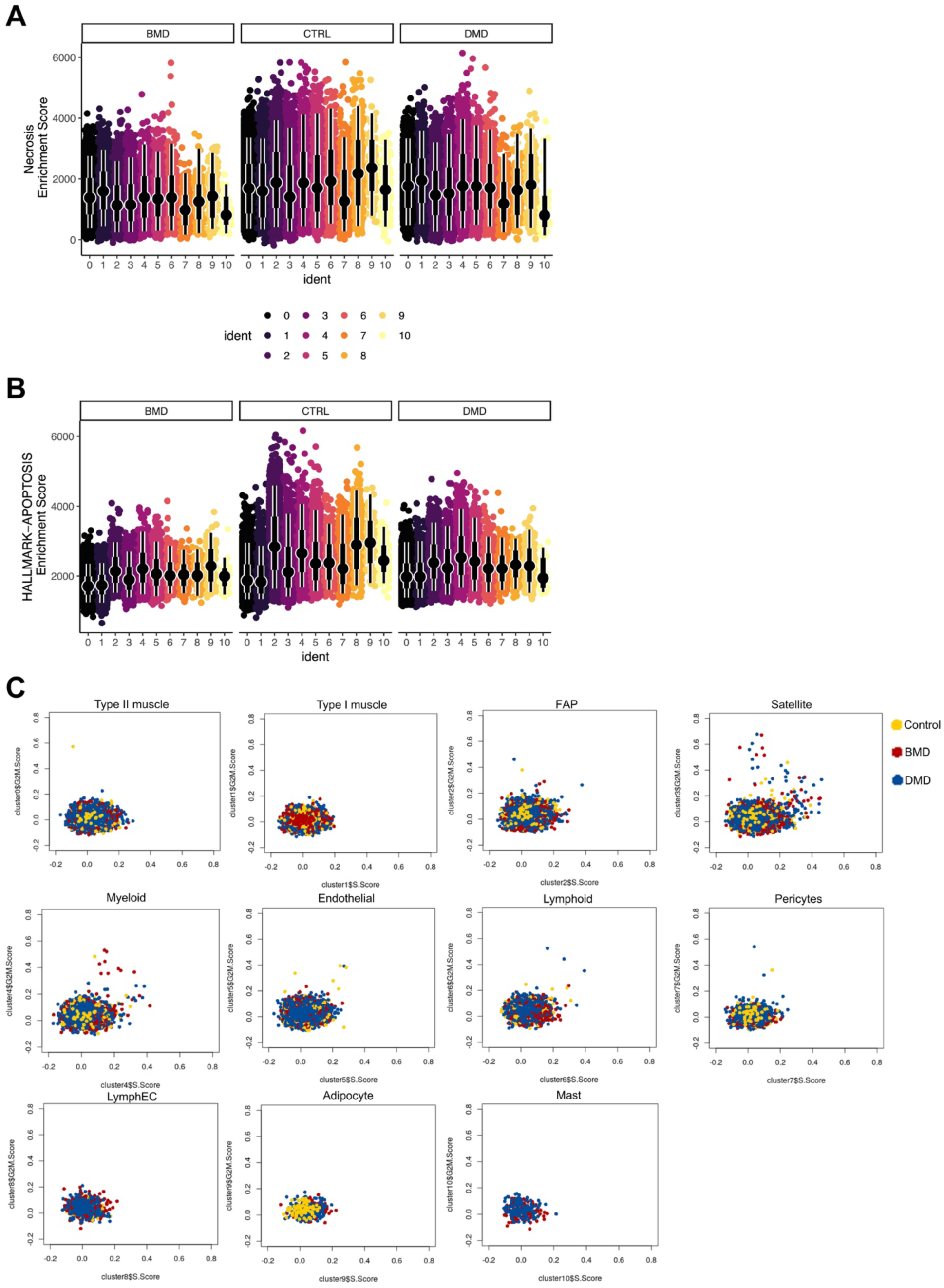
Cell death analysis. **(A)** Plot of necrosis (GSEA systematic name M41803) enrichment score. (**B**) Plot of apoptosis (GSEA systematic name M5902) enrichment score. (**C**) Scatterplot of cell cycle scores in all cell types. Numeric labels denote different cell types: 0 - Type II muscle, 1 - Type I muscle, 2 - Fibro-adipogenic progenitors (FAP), 3 - Satellite cells, 4 - Myeloid cells, 5 - Endothelial cells (EC), 6 - Lymphoid cells, 7 - Pericytes, 8 - Lymphatic endothelial cells (lymphEC), 9 - Adipocytes, 10 - Mast cells.

**Fig. S7.**
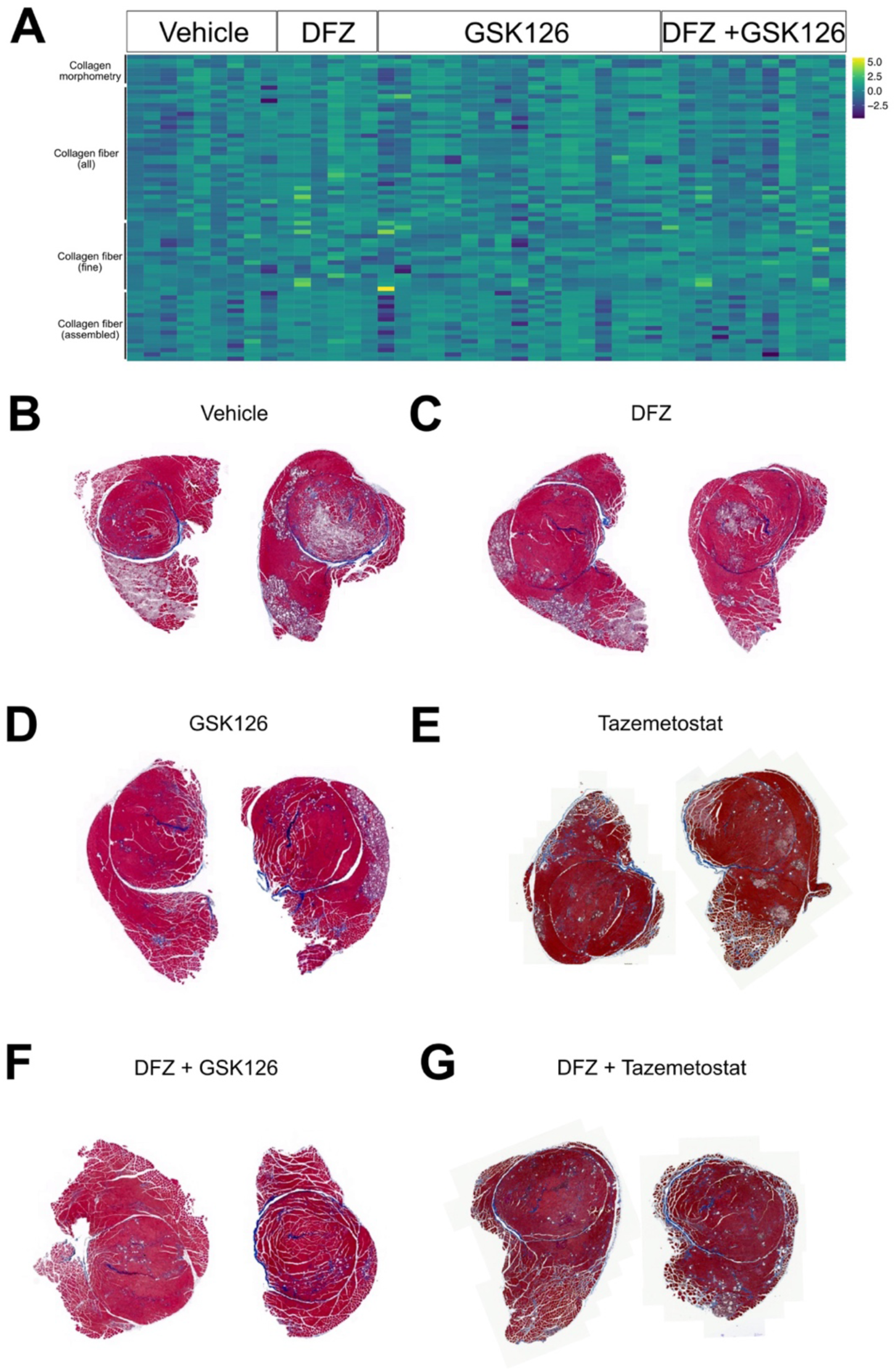
Image analysis of Masson’s trichrome staining. **(A)** Heatmap illustrating the quantification of images for collagen morphometry, collagen fiber, fine collagen fiber, and assembled fiber. (**B-G**) Representative images of Masson’s trichrome staining for the vehicle group (**B**), the deflazacort-injected group (**C**), the GSK126-injected group (**D**), the Tazemetostat-injected group (**E**), the deflazacort and GSK126-injected group (**F**), and the deflazacort and Tazemetostat-injected group (**G**).

**Fig. S8.**
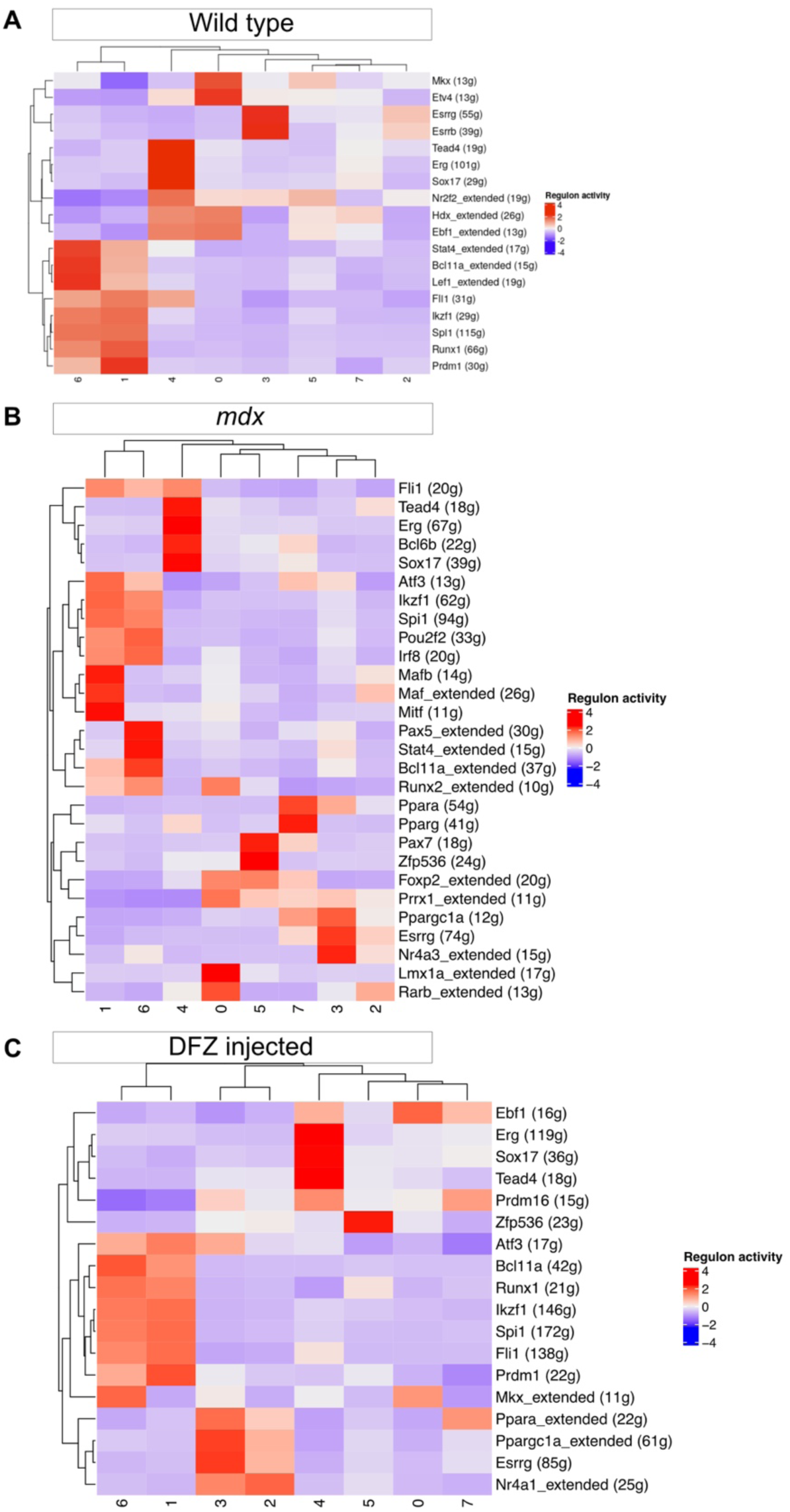
Analysis of ligand-receptor interactions. **(A)** Heatmap of significant ligand-receptor pairs in WT. (**B**) Heatmap of significant ligand-receptor pairs in *mdx* mice. (**C**) Heatmap of significant ligand-receptor pairs in *mdx* with deflazacort injection. Numeric labels denote different cell types: 0 - Type II muscle, 1 - Type I muscle, 2 - Fibro-adipogenic progenitors (FAP), 3 - Satellite cells, 4 - Myeloid cells, 5 - Endothelial cells (EC), 6 - Lymphoid cells, 7 - Pericytes, 8 - Lymphatic endothelial cells (lymphEC), 9 - Adipocytes, 10 - Mast cells.

**Fig. S9.**
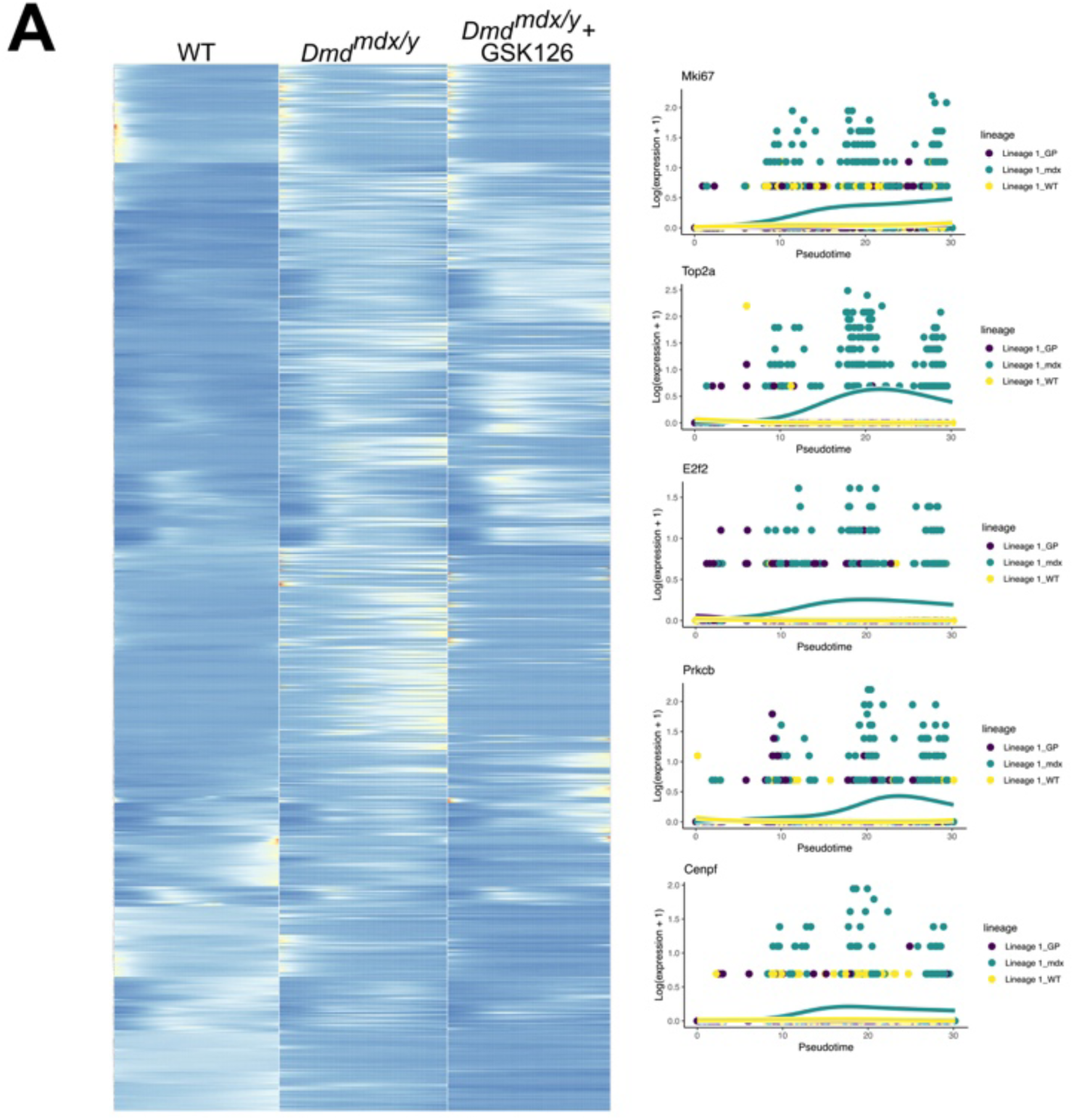
Heatmap of pseudotemporal ordering of satellite and muscle cells along a path towards a differentiated state. **(A)** Heatmap and plot of log-transformed counts and the fitted values of wild type (WT), *D2-mdx*, *D2-mdx* with GSK126 (GP) injection.

**Fig. S10.**
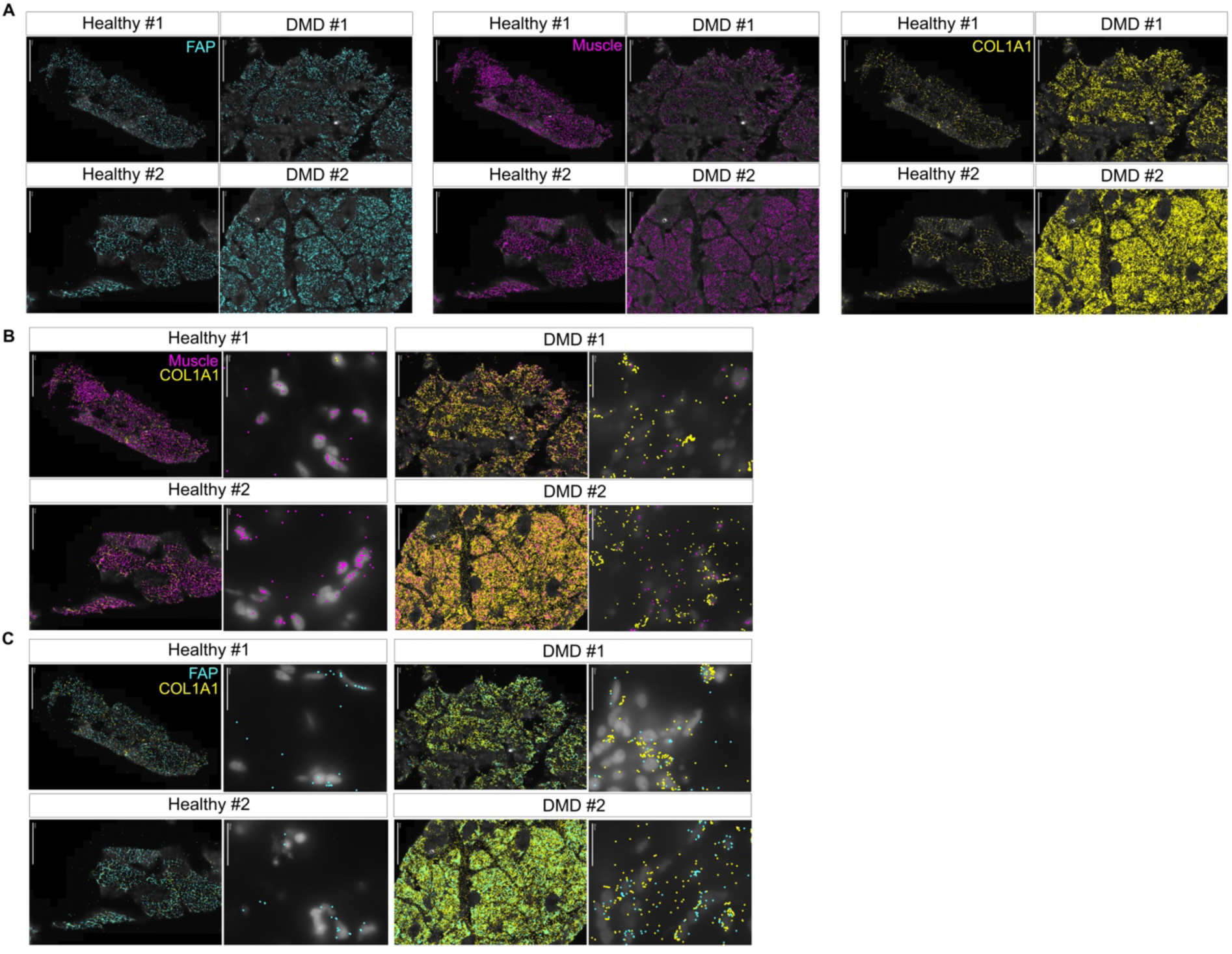
Spatial map of human muscle tissues using MERFISH. **(A)** Spatial expression of FAP (*CCDC102B, COL4A1, COL4A2, LAMA4, NRXN1*) markers, muscle markers (*ANKRD1, ERBIN, PFKFB3, PLIN5, TTN, WNK2, XIRP2*), and fibrosis marker *COL1A1*. (**B**) Co-expression map of muscle markers and fibrosis marker. (**C**) Co-expression map of FAP and fibrosis marker.

**Fig. S11.**
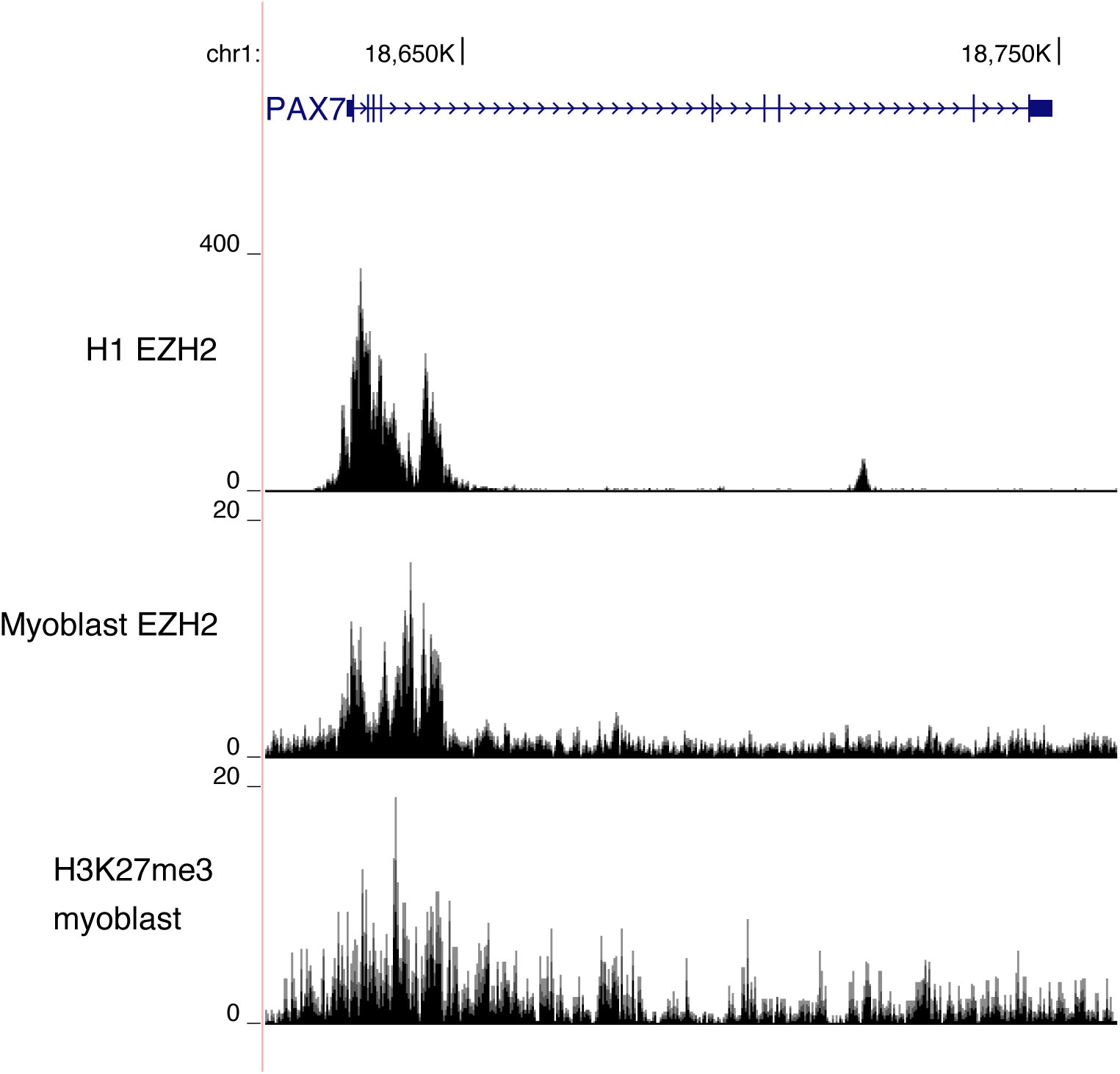
EZH2 ChIP-seq peak locus on *PAX7*.

**Fig. S12.**
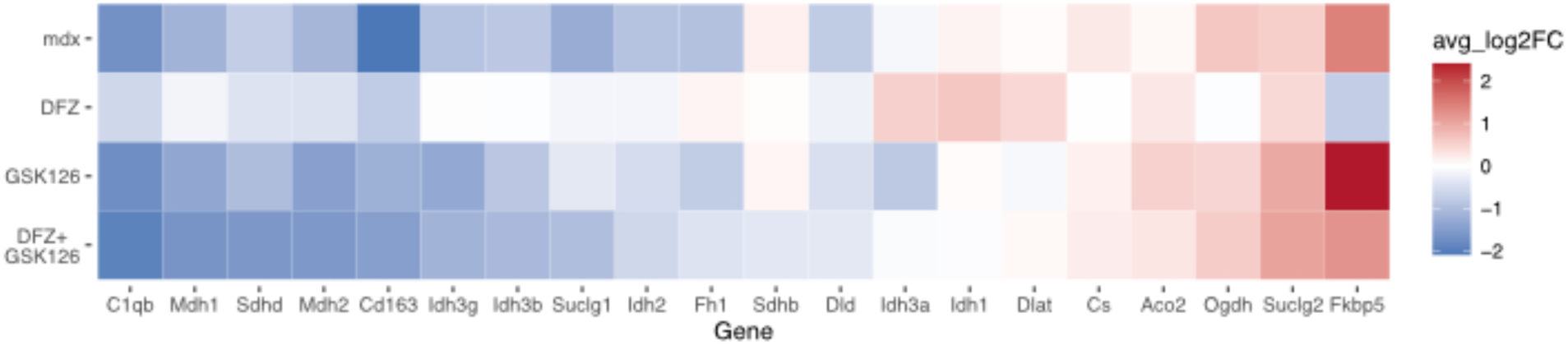
Heatmap of gene expression patterns related to TCA cycle in myeloid cells by the addition of deflazacort and EZH2 inhibitor.

**Table S1.** Information of study participants.

| Sample | Sex | Age | Mutation | Steroid treatment | Biopsy area | Used for Visium/MERFISH analysis |
| --- | --- | --- | --- | --- | --- | --- |
| Healthy |  |  |  |  |  |  |
| 1 | M | 16 | - | No | Quad | Yes/Yes |
| Healthy |  |  |  |  |  |  |
| 2 | M | 3 | - | No | Quad | No/Yes |
| Healthy |  |  |  |  |  |  |
| 3 | M | 3 | - | No | Abdomen | No/No |
| Healthy |  |  |  |  |  |  |
| 4 | F | 5 | - | No | Abdomen | No/No |
| Healthy |  |  |  |  |  |  |
| 5 | M | 10 | - | No | Abdomen | No/No |
| BMD1 | M | 13 | p.Ser1273X | No | Quad | No/No |
|  |  |  | p.Glu2147_Gln2171d |  |  |  |
| BMD2 | M | 4 | el | No | Quad | No/No |
| BMD3 | M | 7 | exon 30-42 deletion | No | Quad | No/No |
| DMD1 | M | 17 | p.Arg195X | No | Quad | No/No |
| DMD2 | M | 5 | p.Gln423* | No | Quad | Yes/Yes |
| DMD3 | M | 5 | p.His3299Glnfs*15 | No | Quad | Yes/Yes |

**Table S2.** Genes denoting each cell type.

| Organism | Cell type | Marker gene used |
| --- | --- | --- |
| Human | Type II muscles | MYH1, MYH2 |
| Human | Type I muscles | MYH7 |
| Human | FAP | PDGFRA |
| Human | Satellite | PAX7 |
|  | Immune cells | PTPRC, CD14, CD163, C1QA, |
| Human | (myeloid) | S100A8 |
| Human | EC | PECAM1 |
|  | Immune cells |  |
| Human | (lymphoid) | IL7R, PTPRC, NKG7, PRF1 |
| Human | Pericyte | PDGFRB |
| Human | LymphEC | TFF3 |
| Human | Adipocyte | PLIN1, ADIPOQ |
| Human | Mast cells | TPSB2 |
| Mouse | FAP | Pdgfra |
| Mouse | Myofiber | Myh1, Myh2, Myh7 |
|  | Immune cells |  |
| Mouse | (myeloid) | Ptprc, Cd14, F13a1, Clec10a |
|  | Immune cells |  |
| Mouse | (lymphoid) | Ptprc, Cd79a |
| Mouse | EC | Pecam1 |
| Mouse | Satellite | Pax7 |
| Mouse | Adipocyte | Adipoq |

**Data S1. Differentially expressed genes in human and mouse (separate file).**

**Data S2. FibroNest analysis terms (separate file).**

